# AdaptivePy: a unified Python framework for adaptive sampling in molecular dynamics

**DOI:** 10.64898/2026.08.04.742868

**Authors:** Hassan Nadeem, Diego E. Kleiman, Diwakar Shukla

**Author notes:** These authors contributed equally to this work (Hassan Nadeem, Diego E. Kleiman).

## Abstract

Adaptive sampling accelerates the exploration of conformational space in molecular dynamics (MD) simulations by repeatedly analyzing the accumulated trajectories and seeding a new round of simulations from informative configurations. A growing collection of adaptive sampling policies has been proposed, each built around a particular notion of what makes a configuration informative, yet these methods are scattered across separate and often incompatible implementations, which complicates their systematic comparison and their combined use in meta adaptive sampling schemes. Here, we present AdaptivePy, a compact and extensible Python framework that implements nine seed-selection policies behind a single configuration-driven interface, spanning simple population-based baselines, several established machine-learning and geometry-based methods, and two ensemble or meta sampling policies introduced in this work. We show that the shared implementation reproduces the characteristic selection behavior of each policy on a series of analytic benchmark landscapes. We also introduce a new adaptive sampling scheme that employs TS-DAR, a deep learning framework originally designed to identify transition states, into an acquisition criterion that drives the discovery of an entire multi-basin landscape starting from a single basin. We further demonstrate that the common interface enables meta adaptive sampling policies, which aggregate the rankings of several policies into a single set of seeds. AdaptivePy thereby provides a unified testbed for the adoption, benchmarking, and continued development of adaptive sampling methods for biomolecular MD simulations.

## Introduction

Machine-learning (ML) methods have substantially advanced the prediction of protein structure, ^1–3^ conformational heterogeneity, ^4–7^ dynamics, ^8–11^ and ensemble properties. ^12–15^ Nevertheless, physics-based molecular dynamics (MD) simulations remain essential because they provide explicit atomistic dynamics generated from defined potential-energy functions and, under appropriate conditions, can be used to estimate thermodynamic ^16^ and kinetic observables. ^17^ By contrast, many current ML-based generative models are not designed to reproduce equilibrium distributions or dynamical statistics with physical fidelity. ^18^

The numerical stability of conventional MD integrators requires time steps on the order of femtoseconds. Consequently, the duration of an individual continuous trajectory often remains substantially shorter than the biologically relevant timescales associated with processes such as signal transduction, ^19–21^ enzymatic catalysis, ^22,23^ protein folding, ^24,25^ solute transport, ^26,27^ ligand (un)binding ^28,29^ and large-scale conformational rearrangements. ^30–32^ This disparity between accessible simulation timescales and the timescales of molecular events has motivated the development of numerous enhanced-sampling strategies. ^33^

Enhanced-sampling methods can be broadly classified according to how they accelerate exploration of configuration space. ^33^ One class modifies the sampled distribution or underlying dynamics by introducing biasing potentials, ^34^ selectively accelerating particular degrees of freedom, ^35^ or varying thermodynamic parameters. ^36^ A second broad class preserves unbiased trajectory segments but redistributes computational effort at the trajectory level. Adaptive sampling methods do so by selecting previously sampled configurations as starting points for additional simulations, ^37,38^ whereas related trajectory-stratification approaches include milestoning ^39^ and weighted ensemble. ^40^

Adaptive sampling methods iteratively analyze accumulated simulation data and select new starting configurations (seeds) according to an acquisition or ranking criterion. By preferentially initiating trajectories from undersampled, kinetically informative, or otherwise relevant regions of configuration space, these methods can improve conformational exploration and accelerate the convergence of thermodynamic or kinetic estimates relative to non-adaptive simulation strategies. ^41^ The criteria used to identify informative configurations, however, differ substantially among policies. Population-based methods such as least-counts sampling prioritize sparsely populated clusters and therefore provide a simple, computationally inexpensive means of promoting exploration, although their rankings do not account for the geometry, kinetics, or functional relevance of the sampled states. ^37,42^ Geometry-based approaches such as kNN-AS instead identify configurations near the boundaries of the sampled region from their local neighborhood structure, allowing them to target exploratory frontiers without requiring a predefined reaction coordinate, but making their behavior dependent on the chosen feature representation and distance metric. ^43^ Objective-directed methods such as FAST balance exploration of undersampled states with exploitation of user-specified structural features, which can efficiently drive sampling toward a known target but requires an informative objective to be defined in advance. ^44^ MA-REAP similarly learns collective-variable weights from accumulated trajectories and coordinates exploration among multiple simulation agents, providing an adaptive alternative to fixed objective weights at the cost of additional optimization and hyperparameter choices. ^45,46^

Other policies use learned kinetic representations to prioritize configurations that may be especially informative about slow transitions. MaxEnt VAMPNet trains a VAMPNet on time-lagged data and selects frames whose soft metastable-state assignments have high entropy, thereby favoring configurations that cannot be assigned confidently to a single state. ^47,48^ TS-DAR likewise learns a time-lagged representation but ranks frames according to their distance from learned metastable-state prototypes, identifying transition-state-like or out-of-distribution configurations that may connect neighboring basins. ^49^ These learned criteria can incorporate kinetic information that is absent from purely population- or geometry-based policies, but their performance depends on the quality and stability of the trained representation and generally entails greater computational and methodological complexity. More broadly, the effectiveness of any policy depends on the topology and kinetics of the conformational landscape, the representation used to characterize molecular configurations, the amount of accumulated data, and whether the objective is broad exploration, discovery of transitions, or convergence of a particular thermodynamic or kinetic quantity. Consequently, the advantages of one policy can become disadvantages under a different landscape or performance metric. A systematic comparison has shown that no single adaptive sampling policy performs best across all systems and objectives, and that combining complementary policies can provide more robust performance across measures of conformational exploration and statistical convergence. ^41^

To facilitate the systematic application and comparison of adaptive sampling strategies, we developed AdaptivePy, a compact Python package that implements nine conformational seed ranking policies. These comprise two baseline approaches (random selection and least-counts sampling^37^), five advanced methods (FAST, ^44^ MA-REAP, ^46^ kNN-AS, ^43^ MaxEnt VAMPNet, ^48^ and TS-DAR, ^49^ an existing MD analysis method adapted here for use as an adaptive sampling criterion), and two meta-policies (allocation and majority polling). We evaluate AdaptivePy across several molecular systems, reproduce algorithm behavior, and demonstrate methodological combinations enabled by implementing these approaches within a common framework. Beyond the policies currently implemented, numerous adaptive sampling strategies have been proposed, including AdaptiveBandit ^50^ and TSLC. ^51^ Owing to its modular and extensible design, AdaptivePy readily supports the implementation and integration of these methods, as well as future adaptive sampling policies, within a unified framework. While previous studies had implemented software to carry out adaptive sampling simulations under specific conditions or computational environments, ^52,53^ we anticipate that our general, multi-policy package will support the adoption, benchmarking, and continued development of adaptive sampling methods for biomolecular MD simulations.

## Results and discussion

### AdaptivePy is an extensible framework for adaptive sampling MD simulations

AdaptivePy organizes adaptive sampling as an explicit iterative loop wrapped around an existing simulation engine (Figure 1a). At the beginning of each new round of simulations, the trajectories accumulated up to last round are reduced to featurized trajectories. These are numerical arrays whose columns may be any structural or collective coordinates that the user considers relevant to the process under study. For instance, root-mean-square deviations, solvent-accessible surface areas, inter-residue distances, or lower-dimensional collective variables have been used as collective variables for adaptive sampling. These accumulated features are then optionally partitioned by AdaptivePy into clusters that discretize the region of feature space explored so far. A seed-selection policy assigns an acquisition score to each cluster, or to each individual frame, and the highest-scoring configurations are returned as the starting structures for the next round of simulation. Iterating this cycle concentrates simulation effort in the regions that the chosen policy judges most informative, without biasing the underlying equations of motion, and keeping the unbiased character of each individual trajectory unchanged.

**Figure 1:**
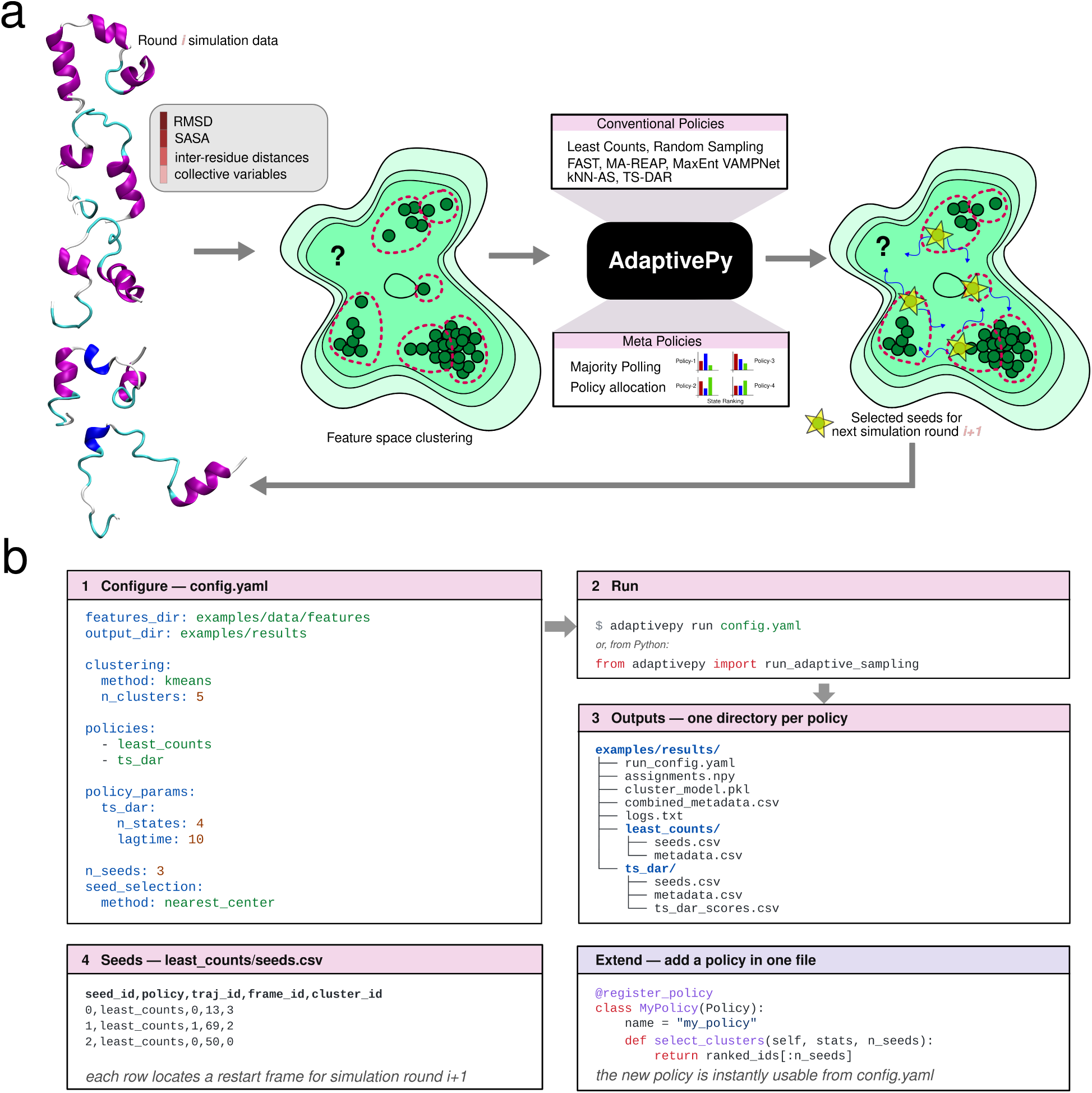
AdaptivePy adaptive sampling workflow and interface. (a) Each simulation round is converted into feature trajectories and divided into clusters. AdaptivePy ranks these clusters using seven conventional policies: least-counts, random sampling, FAST, MA-REAP, MaxEnt VAMPNet, kNN-AS, and TS-DAR. It also supports two meta-policies, majority polling and allocation. The selected configurations seed the next simulation round. (b) Users define the clustering, policies, and seed settings in a YAML file. AdaptivePy runs from the command line or Python and saves the selected seeds, metadata, and clustering results. New policies can also be implemented as Python classes through inheritance and invoked directly from the configuration file.

The policies bundled with AdaptivePy differ chiefly in how they translate accumulated data into an acquisition score. The framework is deliberately structured so that this translation is the only component a new method needs to supply. Most policies act at the cluster level and rank clusters by their population, their geometry within feature space, or a learned reward. The two deep learning policies, MaxEnt VAMPNet and TS-DAR, instead operate at the level of individual frames: each trains a model on time-lagged pairs of feature vectors and scores frames directly, so that clustering becomes optional and is imposed only when these methods are combined with cluster-level policies. When a cluster is selected, the representative seed frame is drawn from it according to a configurable rule, taking either the frame closest to the cluster center or a uniformly random frame from the cluster. A single human-readable configuration file specifies the feature inputs, the clustering scheme, the policy and its parameters, the number of seeds requested, and the seed-selection rule, so that an entire round is reproducible from one text file, (Figure 1b). A new policy can be introduced by implementing a compact scoring routine that reuses the shared clustering, cluster-statistics, and input and output machinery. A further consequence of this common structure is that AdaptivePy can run several policies concurrently on the same accumulated data and reconcile their separate rankings into a single set of seeds. Because each policy exposes its decision in the same ranked form, this reconciliation is handled by a lightweight aggregation layer rather than by any modification of the policies themselves, which is what allows the meta-policies examined later in this work to be deployed using the same framework.

### AdaptivePy recapitulates algorithm behavior in benchmark potentials

A framework that reimplements several published algorithms must first establish that each reimplementation is faithful to its original description. Analytic low-dimensional potentials are particularly well suited to this purpose, because their minima, barriers, and relative basin populations are known exactly, so that the configurations a policy chooses to revisit can be interpreted directly rather than inferred from a low-dimensional projection of a high-dimensional system. We therefore generated reference trajectories on a set of two-dimensional model potentials that differ in the number, depth, and connectivity of their basins. These comprise the Müller–Brown surface, a symmetric four-well potential, and the wave-split, smooth-minima, and triple-well potentials, together with the three-dimensional chaotic Lorenz attractor (see Methods). We then applied a single policy to the pooled trajectories of each system (Figure 2). Because the underlying landscapes are known, the placement of the selected seeds can be read directly against the objective that each method was designed to optimize.

**Figure 2:**
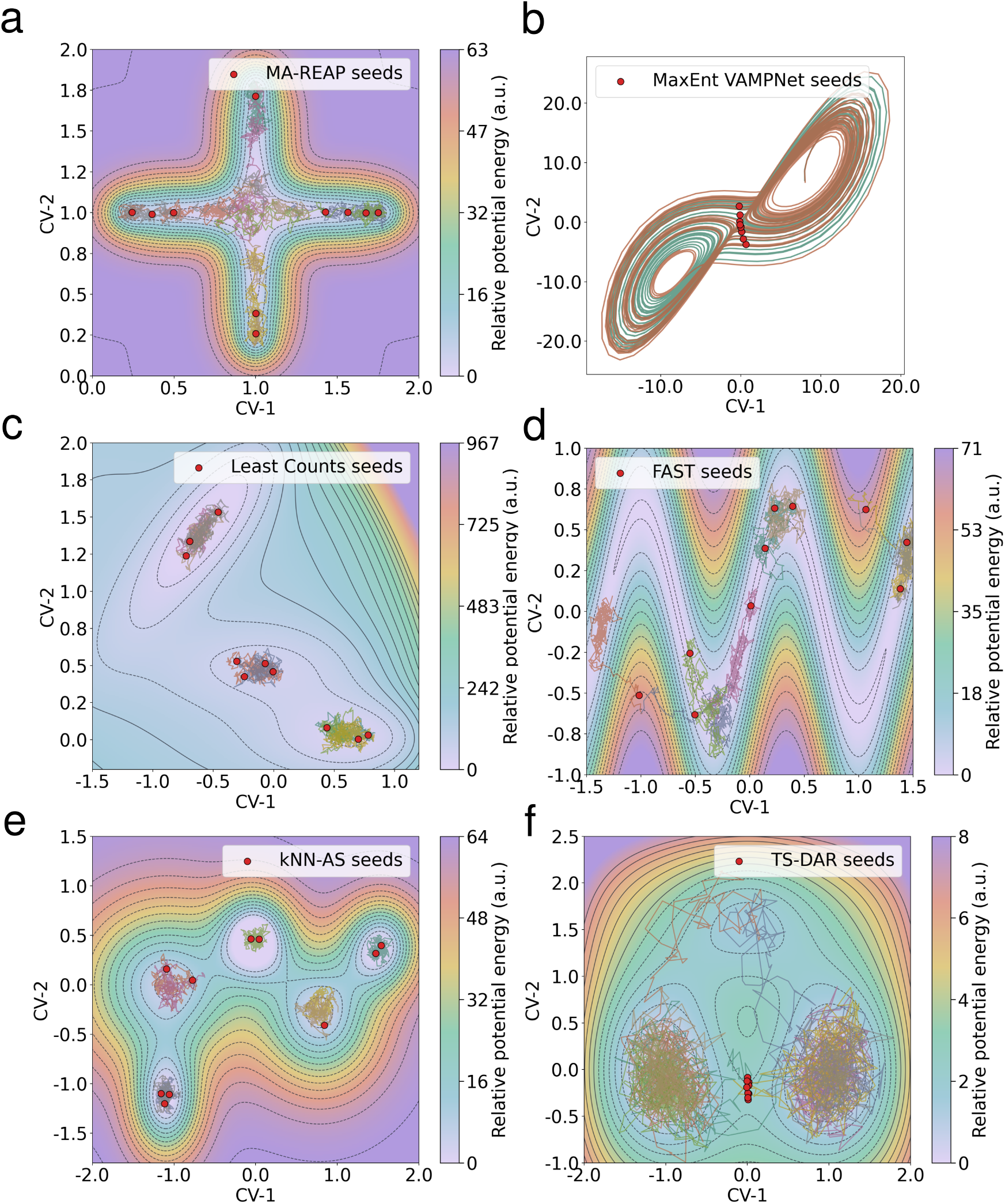
AdaptivePy reproduces the characteristic seed-selection behavior of each policy on benchmark systems. Seeds selected in a single round (red) are overlaid on the sampled trajectories and the underlying energy landscape. Each system was sampled with 8 trajectories of 3500 frames (random seed 42); ten seeds were selected per system. Cluster-level policies (a, c, d, e) used 20 *k*-means clusters, whereas the frame-level policies (b, f) scored frames directly. (a) MA-REAP on the symmetric four-well potential. (b) MaxEnt VAMPNet on the chaotic Lorenz attractor. (c) Least-counts sampling on the Müller–Brown potential. (d) FAST, configured to maximize both coordinates, on the wave-split potential. (e) kNN-AS on the smooth-minima potential. (f) TS-DAR on the triple-well potential. Color bars give the relative potential energy in arbitrary units; axes are the two feature coordinates (CV-1, CV-2).

Each policy produced the selection pattern anticipated from its formulation. MA-REAP, which partitions the trajectories among agents that learn separate collective-variable weightings, distributed its seeds along both arms of the four-well cross, following the two orthogonal directions its agents had identified as informative and thereby coordinating rather than duplicating their exploration (Figure 2a). MaxEnt VAMPNet placed its seeds within the narrow crossover connecting the two lobes of the Lorenz attractor, where the soft assignment of frames to metastable states is most ambiguous and the Shannon entropy of that assignment is largest (Figure 2b). Least-counts sampling, the simplest of the exploration baselines, spread its seeds across the sparsely populated clusters of the Müller–Brown surface and away from the trajectory-dense interiors of its basins (Figure 2c). FAST, configured to maximize both coordinates, biased its seeds toward the high-coordinate region of the wave-split potential while retaining a fraction in undersampled clusters, a direct expression of the balance between exploitation and exploration that lies at the center of the method (Figure 2d). kNN-AS selected seeds in the outer, boundary basins of the smooth-minima landscape, consistent with its use of a nearest-neighbor graph to identify states that sit on the frontier of the sampled region (Figure 2e). Finally, TS-DAR concentrated its seeds at the saddle separating the two lower basins of the triple-well potential, the transition region that its out-of-distribution score is constructed to flag (Figure 2f). Reproducing six qualitatively different selection strategies, namely directed, entropy-based, population-based, geometric, and barrier-seeking exploration, from a single codebase indicates that the shared implementation preserves the distinct intent of each algorithm. Beyond confirming correctness, this consistency is what makes the framework useful as a benchmarking platform: because every method consumes identical inputs and is configured through the same interface, subsequent differences in their behavior can be attributed to the acquisition criteria themselves rather than to incidental differences in implementation, featurization, or preprocessing.

### Demonstrating the extensibility of AdaptivePy through TS-DAR

As described previously, AdaptivePy isolates the acquisition function from the rest of the adaptive sampling workflow. This allows new sampling strategies to be introduced by implementing a minimal policy-specific scoring routine. To demonstrate this extensibility, we incorporate TS-DAR. ^49^ The policies considered so far were all originally conceived as sampling methods. TS-DAR, in contrast, was developed as an analysis method: given a set of trajectories that have already been generated, it locates the transition-state conformations that connect the metastable states of a system. ^49^ It accomplishes this by embedding each frame onto a hypersphere in a learned latent space, using a VAMP-2 objective to compress the frames belonging to a common metastable basin toward a shared prototype, and a dispersion regularizer to drive the prototypes of distinct basins apart on the hypersphere. Frames that lie on the barriers between basins are consequently pushed away from every prototype and are flagged as out-of-distribution (OOD) relative to the densely populated basins. We reasoned that this OOD score is also a natural acquisition criterion for sampling. Configurations with a high OOD score occupy the edges of the sampled basins and the barriers between them, which are precisely the points from which a newly launched trajectory is most likely to commit into an adjacent, and possibly still undiscovered, basin. We therefore exposed the per-frame OOD score as a ranking policy and asked whether a method built to describe transition states after a landscape has been sampled could instead be inverted to drive that sampling.

To examine this, we ran the policy as a closed loop on the triple-well potential under a deliberately unfavorable initial condition, launching all ten trajectories of the zeroth round from within a single basin. In each subsequent round the model was retrained on all features accumulated up to that point, the ten frames with the highest OOD score were selected, and ten fresh trajectories of one thousand frames each were initiated from them; the loop was iterated for thirty rounds, and the entire protocol was repeated for six independent random seeds. The cumulative sampling plots of a representative run trace how the sampled region grows outward from the starting basin over these rounds (Figure 3a–d). After the first round the trajectories remain essentially confined to the initial well; by the tenth round the loop has crossed the low barrier into the second deep basin and has begun to ascend toward the shallower upper basin; and by the twentieth and thirtieth rounds all three basins, together with the elevated rim surrounding them, have been populated.

**Figure 3:**
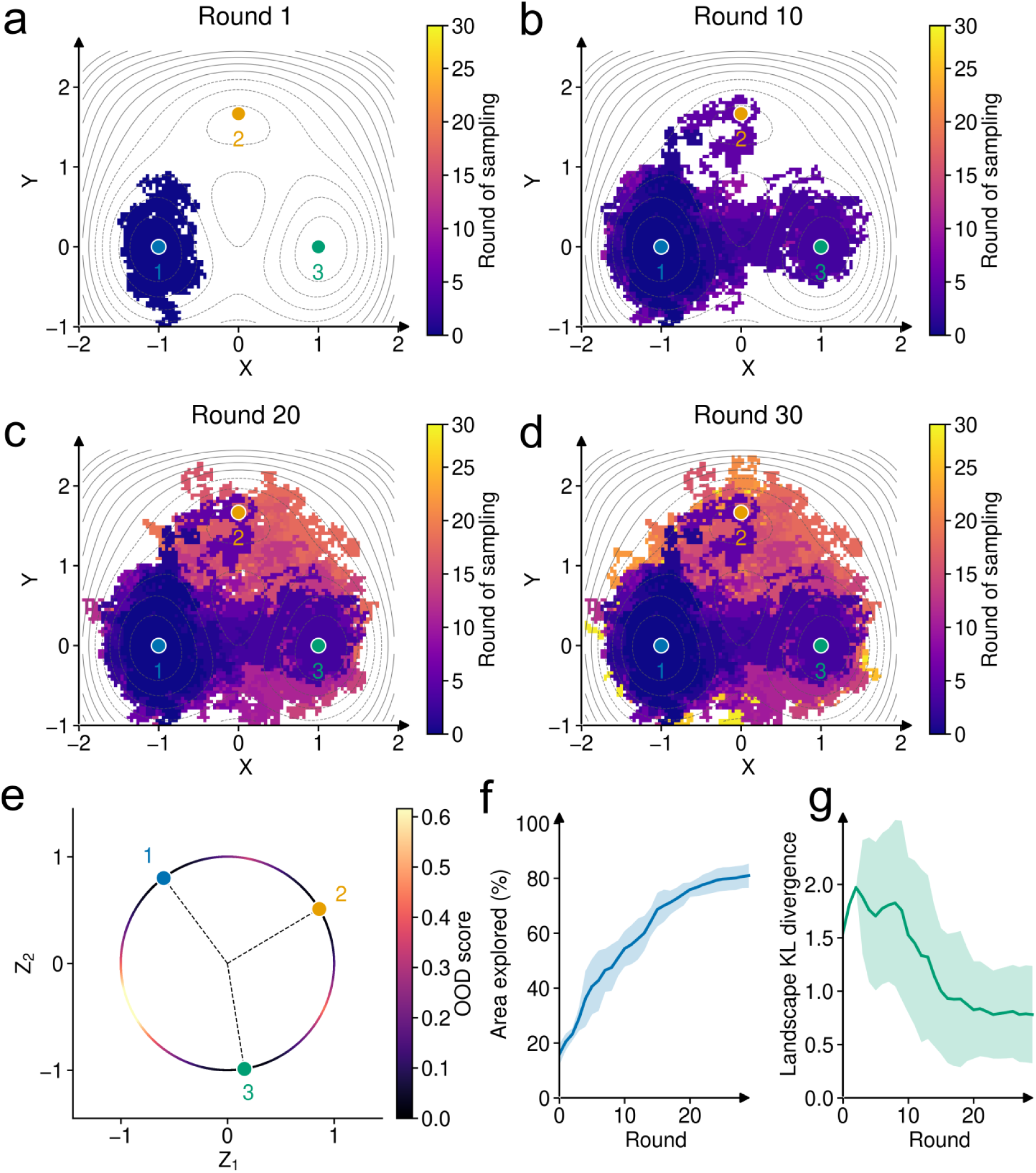
TS-DAR out-of-distribution scores drive adaptive sampling of the triple-well potential. All round-zero trajectories were initialized in a single basin. Each round retrained TS-DAR on the accumulated features, selected the ten highest out-of-distribution (OOD) frames, and restarted ten trajectories of 1000 frames from them, for 30 rounds. (a–d) Sampling maps for a representative run at rounds 1, 10, 20, and 30, colored by the round in which each region was first sampled; gray contours are the reference free energy and the numbered points mark the three basin minima. (e) All sampled frames re-embedded through the final-round model onto the unit circle, colored by OOD score; the three basins form well-separated prototypes (points) and the OOD score is largest between them. (f) Fraction of the landscape that has been visited and (g) Kullback–Leibler divergence between the sampled and reference Boltzmann distributions, as a function of round. Lines and shaded bands are the mean ± one standard deviation over six random seeds.

Re-embedding every sampled frame through the final trained model confirms the mechanism responsible for this expansion (Figure 3e). The three basins map onto well-separated prototypes distributed around the unit circle, and the OOD score increases smoothly along the arcs between them, so that the frames ranked most highly by the policy are exactly those that bridge neighboring metastable states. The exploration is summarized by two convergence measures averaged over the six random seeds. The fraction of the landscape that had been visited rose from 16 ± 4% at initialization to 81 ± 4% after thirty rounds (Figure 3f). The Kullback–Leibler divergence between the sampled distribution and the reference Boltzmann distribution increased over the first several rounds, reaching a value near 2 as the loop briefly oversampled the starting basin before escaping it, and then decayed to 0.78 ± 0.46 as the remaining basins filled in (Figure 3g). The relatively wide spread of the final divergence is attributable almost entirely to a single run whose network collapsed two of the three prototypes and never learned to separate a basin that its trajectories had in fact visited, leaving a distorted population estimate (divergence 1.66); when that run is excluded, the remaining five converge to 0.60 ± 0.17. Taken together, these results show that the TS-DAR adapted policy drives exploration reliably across independent seeds, whereas the fidelity of the recovered equilibrium populations depends on the quality of the learned embedding, which is a property of the model’s training rather than of the sampling strategy itself.

### AdaptivePy enables exploration of meta adaptive sampling policies

The methods surveyed above rank configurations according to different, and at times conflicting, notions of value, and no single criterion can be expected to be optimal for every landscape or every simulation objective; combining complementary policies has previously been found to improve robustness. ^41^ The unified design of AdaptivePy makes such combinations straightforward, because every policy expresses its decision in the same form, as a ranked list of clusters. Several policies can therefore be evaluated on the same accumulated data and their separate rankings reconciled into a single set of seeds. Frame-level policies are placed on the same footing as cluster-level ones by assigning each cluster the maximum score attained by any of its member frames, so that entropy- and OOD-based methods can be pooled with population- and geometry-based methods without special handling. On this shared representation we implemented two meta-policies that embody two distinct philosophies of aggregation (Figure 4a).

**Figure 4:**
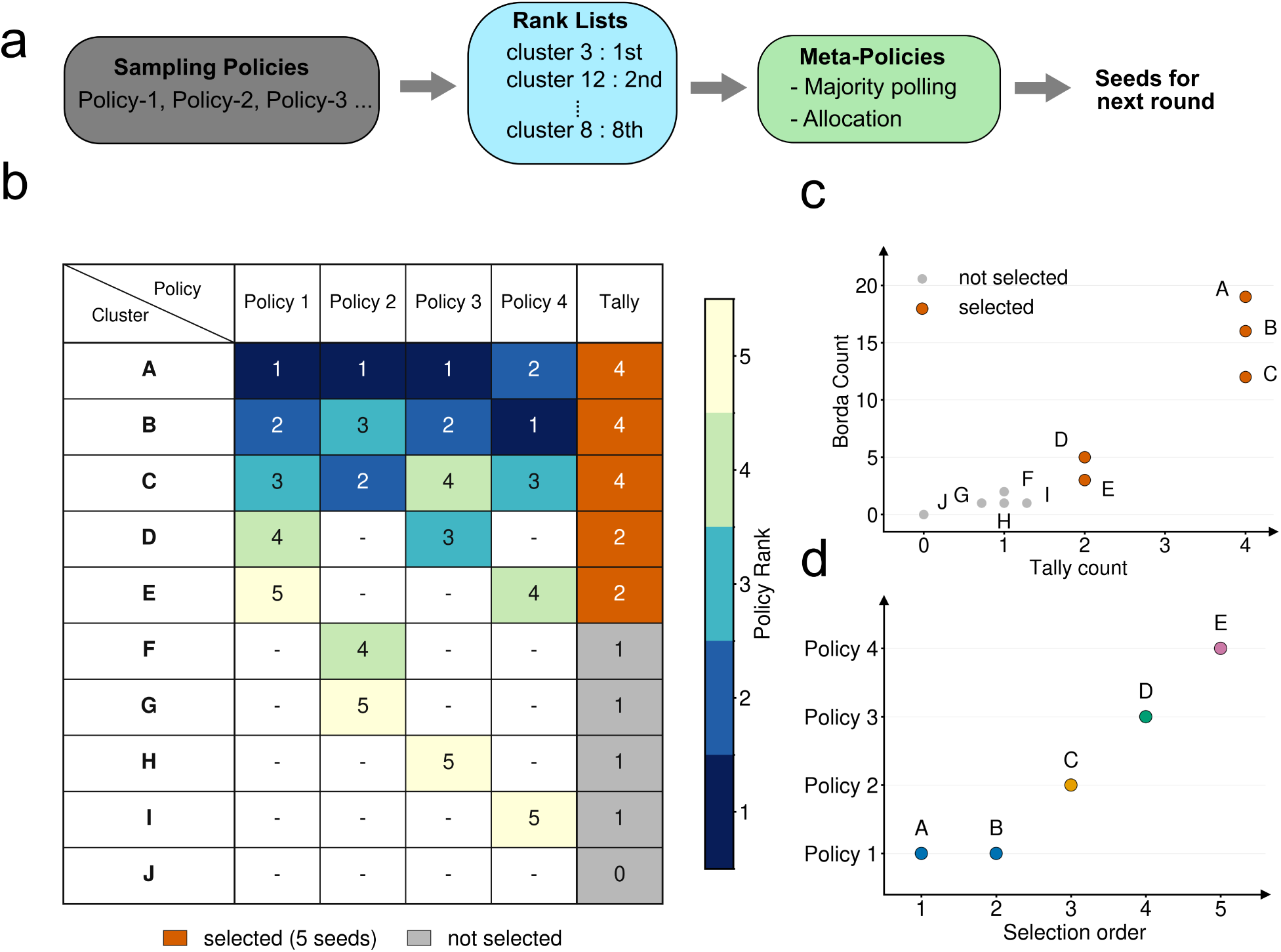
Meta-policies aggregate heterogeneous policies into a single seed set. (a) Each component policy emits a ranked list of clusters, which a meta policy (either majority polling or allocation) combines into the seeds for the next round. (b–d) A worked example with four component policies, ten clusters (A–J), and five seeds. (b) Rank matrix: each policy’s top-five clusters (colored by rank; a dash denotes an unranked cluster), with the per-cluster vote tally in the right-hand column and the majority-polling selection shaded orange. (c) Majority polling orders clusters by tally count and breaks ties by Borda count, selecting the five highest ranked clusters. (d) Allocation instead fills a fixed per-policy quota in policy order, skipping clusters that have already been claimed; labels give the cluster taken at each selection step.

The first, majority polling, selects by consensus. Each participating policy contributes a fixed number of its most highly ranked clusters; the number of policies that rank a given cluster then defines its vote tally, while a Borda-style score sums the rank-weighted points that the cluster accumulates across those policies, so that a cluster placed near the top of several lists scores more than one that only barely qualifies on each. Clusters are ordered first by their tally and only then by their Borda score (Figure 4b,c). Because the tally is the primary key, a cluster endorsed by more policies always outranks one endorsed by fewer, and the Borda score serves to separate clusters that are tied on the tally. User-assigned weights, which give more trusted policies a larger say, enter only through the Borda score and thus act solely as a tie-breaker among equally endorsed clusters; they cannot overturn the consensus itself. The second meta-policy, allocation, is procedural rather than consensual: it visits the policies in a fixed order, grants each a predetermined quota of seeds drawn from the top of its own ranked list, and skips any cluster that an earlier policy has already claimed (Figure 4d). Whereas majority polling can allow a strong consensus among several policies to crowd out a method that finds itself in the minority, allocation guarantees that every participating policy contributes seeds to the next round, preserving the diversity of exploration strategies by construction. The two meta-policies thus trade off against each other: majority polling concentrates seeds where independent criteria agree, whereas allocation deliberately distributes them across criteria even where those criteria dissent.

These strategies are useful only to the extent that the underlying policies genuinely disagree about which configurations to prioritize, and in practice they frequently do, so the aggregation step performs substantive work rather than ratifying a decision the policies had already reached in common. Computationally, the additional cost of ranking the clusters and combining their votes remains small relative to the analysis already performed by the individual component policies.

## Conclusions

AdaptivePy provides a user-friendly and extensible framework that consolidates multiple adaptive sampling algorithms within a common software architecture. Although these methods differ in their acquisition criteria, they share several computational steps, including trajectory featurization, clustering, evaluation of cluster populations, and calculation of distances or related quantities in feature space. AdaptivePy exploits this common structure to provide an efficient implementation in which alternative policies can rank candidate seeds using consistent inputs and configuration parameters.

Because the implemented policies operate on numerical feature trajectories rather than directly on simulation engine-specific data structures, AdaptivePy remains agnostic to both the molecular simulation software and the upstream trajectory analysis pipeline. This abstraction allows the package to be integrated into a range of existing workflows while exposing its functionality through concise configuration files. Moreover, the computational cost of seed selection is generally small relative to the expense of generating atomistic trajectories for biomolecular systems, such that adaptive seed selection introduces limited overhead into the overall simulation workflow.

Implementing these methods within a unified framework also enables the construction of meta-policies that combine seeds proposed by multiple algorithms. Such approaches can distribute computational effort across configurations selected according to complementary exploration, exploitation, geometric, or kinetic criteria, potentially improving robustness when no single policy is optimal for a given conformational landscape or simulation objective.

Future development could extend AdaptivePy to support the automated estimation and parameterization of Markovian ^54^ and memory-dependent^55^ kinetic models. These capabilities would facilitate the integration of additional adaptive sampling strategies, including lambda sampling ^56^ and Ensemble Adaptive Sampling Scheme (EASE). ^41^ In the current implementation, kinetic-model construction is delegated to established external software packages, ^57^ allowing AdaptivePy to remain focused on the standardized ranking and selection of starting configurations.

## Methods

### Langevin dynamics

Two-dimensional benchmark trajectories were generated by integrating overdamped Langevin dynamics on analytic potential-energy surfaces. For each potential, an ensemble of independent trajectories was initialized near the known basin minima. Initial positions were assigned cyclically across the minima, randomly permuted across trajectories, and perturbed by Gaussian noise with standard deviation 10^−3^ in each coordinate. Positions were then propagated with an Euler–Maruyama update,

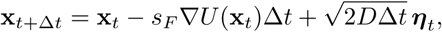

where *U* (**x**) is the analytic potential, *s_F_* is a force-scaling factor, *D* is the diffusion coefficient, and ***η****_t_* ∼ *N* (**0**, **I**) (standard Gaussian noise). Reflecting boundary conditions were applied after every integration step (coordinates leaving the rectangular domain were reflected through the violated boundary and then clipped to the allowed interval).

For Fig. 2, coordinates were saved every 20 integration steps and used directly as the two-dimensional feature vectors for adaptive sampling. Each system was simulated with 8 independent trajectories, 3500 saved frames per trajectory, an integration time step of 10^−4^, and random seed 42. Potential-specific parameters were *D* = 0.015, *s_F_* = 0.003, and bounds [−1.5, 1.2] × [−0.2, 2.0] for Müller–Brown; ^58^ *D* = 0.015, *s_F_* = 0.006, and bounds [0, 2] × [0, 2] for the symmetric four-well potential; ^46^ *D* = 0.02, *s_F_* = 0.01, and bounds [−1.5, 1.5] × [−1, 1] for wave-split; ^43^ *D* = 0.02, *s_F_* = 0.01, and bounds [−2, 2] × [−1.8, 1.5] for smooth-minima; ^43^ and *D* = 1.09^2^*/*2, *s_F_* = 1.0, and bounds [−2, 2] × [−1, 2.5] for the triple-well potential. ^57^

For Fig. 3, the triple-well potential was simulated with 10 independent trajectories per round, 50 integration steps per frame, 1000 saved frames per trajectory, an integration time step of 10^−5^, and varying random seeds. First round trajectories were only initialized in the minimum near (−1, 0). Other parameters are identical to the previous paragraph.

The benchmark potentials were analytic low-dimensional landscapes. The exact functional forms (including parameters) for every potential are reported in Supporting Methods.

### Lorenz system

Chaotic Lorenz ^59^ trajectories were generated with the deeptime.data.lorenz_system simulator. ^57^ Initial conditions were drawn from a Gaussian distribution around (8, 7, 15) with standard deviation 0.5 in each coordinate; the first trajectory was initialized exactly at (8, 7, 15). 8 trajectories of 3500 frames were generated with random seed 42. The resulting three-dimensional coordinates (*x, y, z*) were saved as feature trajectories and used directly for seed in selection.

### Adaptive sampling and baseline algorithms

Available trajectory frames are represented by feature vectors **x***_n_* ∈ ℝ*^d^*. Cluster-based methods partition the discovered region of feature space into non-overlapping clusters *C* = {*c*_1_*, … , c_M_* }, where *c_i_* is the set of frames assigned to cluster *i*, |*c_i_*| is its cardinality or population size, and **z***_i_* is its representative feature vector (usually the cluster center). A policy assigns each cluster (or each frame for frame-level methods) an acquisition score, 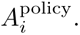 The top *k* ranked clusters or frames are selected as seeds for the next simulation round. For cluster-level policies, the seed frame from a selected cluster is chosen by the configured seed-selection rule, either the frame nearest to the cluster center or a uniformly random frame from the cluster.

Random sampling is used as an undirected baseline. After clustering the discovered feature space, *k* distinct clusters are sampled uniformly at random from *C*, and one seed frame is selected from each sampled cluster.

Least-counts sampling selects the least-populated clusters. ^37^ Let *π* be a permutation of cluster indices such that

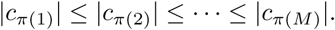

The selected seed clusters are

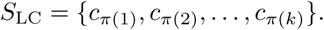

Thus, least-counts is a ranking over clusters, and the top *k* clusters are those with the smallest cardinality or population count.

### FAST

FAST stands for Fluctuation Amplification of Specific Traits.^44^ It is an algorithm that balances directed exploitation of a user-specified feature objective with exploration of undersampled states. For each selected feature *ℓ*, a cluster descriptor *q_iℓ_* is computed as the mean value of feature *ℓ* over frames in cluster *c_i_*. The descriptors are min–max scaled to [0, 1], with the sign chosen according to whether the feature is to be maximized or minimized. The directed score is

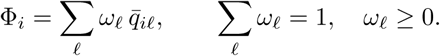

The exploration score is the least-counts component

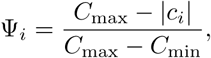

where *C*_min_ and *C*_max_ are the minimum and maximum cluster populations in the current round. The FAST acquisition score is

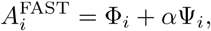

and the *k* clusters with largest 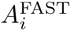 are selected.

### MA-REAP

MA-REAP stands for Multi-agent REinforcement learning based AdaPtive sampling. ^45,46^ This algorithm treats adaptive sampling as a multi-agent allocation problem. Each trajectory is assigned to an agent *a* ∈ {1*, … , N* }. Candidate clusters are first restricted to the *n*_cand_ least-populated clusters. For each candidate *i*, agent stakes *s_ai_* are computed from the number of frames in *c_i_* contributed by agent *a*, using the configured stake rule ^46^ (percentage, equal, max, or logistic; only percentage is used in this work). For each agent, feature means ***µ****_a_*, standard deviations ***σ****_a_*, and non-negative collective-variable weights **w***_a_* are estimated from that agent’s sampled frames. Weights are constrained to sum to one and to change by at most *δ* per adaptive round.

For candidate representative **z***_i_*, the per-agent reward is

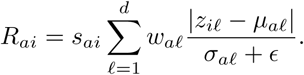

The weights **w***_a_* are optimized by quadratic optimization (namely, SLSQP^60^) to maximize the sum of *R_ai_* over candidate clusters. *ɛ* is a small float added for numerical stability. Per-agent rewards are then combined using the configured MA-REAP regime:

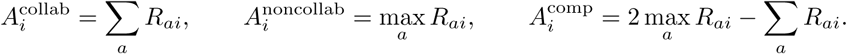

In this study, only the collaborative regime (additive rewards) is used. This parameter was shown to have little to no impact in a previous study. ^46^ Clusters are ranked by the resulting aggregate score, with population and cluster index used as deterministic tie-breakers when necessary.

### kNN-AS

*k*-Nearest-Neighbor Adaptive Sampling (kNN-AS) ^43^ ranks states by local neighborhood geometry in feature space. For each populated cluster, a representative vector **z***_i_* is defined as the cluster center. The *k* nearest neighbors of each representative are identified in Euclidean distance. In the vector-sum mode, the acquisition score is

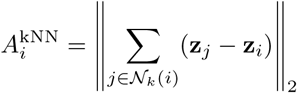

where N*_k_*(*i*) is the set of cluster *i*’s nearest neighbors. In an alternative distance mode, the score is the mean neighbor distance,

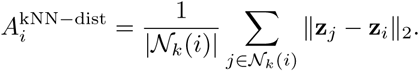

Clusters with the largest kNN-AS scores are selected. Only vector-sum mode is used in this study.

### MaxEnt VAMPNet

MaxEnt VAMPNet is a frame-level policy and therefore does not require clustering (clustering is forced on meta-policies). A VAMPNet ^47^ is trained on lagged trajectory pairs (**x***_t_,* **x***_t_*_+*τ*_) using a duplicated neural-network lobe with softmax output. The model maps each frame to probabilities over *K* soft states,

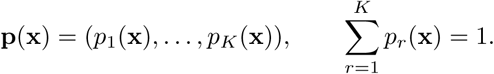

Following the maximum-entropy acquisition criterion, each frame is scored by the Shannon entropy of its soft-state assignment,

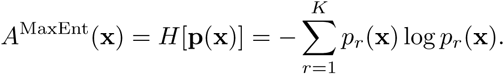

The *k* frames with largest entropy are selected as seeds. Unless otherwise specified, the implementation used lag time *τ* = 1, learning rate 10^−4^, batch size 2048, 100 epochs, numerical regularization *ɛ* = 10^−6^, and hidden-layer widths [128, 64]. The number of softmax states defaulted to the feature dimensionality unless explicitly specified. For the Lorenz example, two states are used.

### TS-DAR

TS-DAR (Transition State identification via Dispersion and vAriational principle Regularized neural networks) ^49^ is also a frame-level policy. A neural network is trained on lagged pairs to learn hyperspherical embeddings **h**(**x**) and soft metastable-state probabilities. The network is optimized with a VAMP-2 objective and, after a pretraining period, a dispersion regularizer that encourages separated state prototypes on the hypersphere. Embeddings are normalized to radius *γ*,

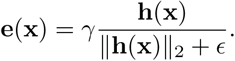

After training, each frame is assigned to its most probable state, and a normalized state center **m***_r_* is computed from the embeddings assigned to state *r*. TS-DAR treats transition-state-like frames as out-of-distribution relative to dense metastable basins. The acquisition score is the cosine distance to the nearest state center,

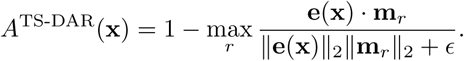

The *k* frames with largest out-of-distribution score are selected. Default parameters are: lag time *τ* = 1, learning rate 10^−3^, batch size 2048, 100 epochs, 10 pretraining epochs, *β* = 0.01, *γ* = 1.0, scaling temperature 0.1, prototype update factor 0.5, Adam optimization, train split 0.9, and *ɛ* = 10^−6^. The default hidden-layer widths are [128, 64].

## Supporting Information

Supporting Information contains functional forms of idealized potentials under Supporting Methods.

## Data and Software Availability Statement

The AdaptivePy Python package, along with the datasets and scripts required to reproduce the figures presented in this work, is publicly available at https://github.com/ShuklaGroup/AdaptivePy. Documentation is available at https://ShuklaGroup.github.io/AdaptivePy/. The package can be installed from PyPI using: pip install adaptivepy-sampling

## Author Contributions

H.N. and D.E.K. contributed equally to this work. H.N., D.E.K., and D.S. designed the study. H.N. and D.E.K. developed the codebase. D.E.K. conducted the experiments and H.N. performed the analyses. H.N. and D.E.K. wrote the manuscript with input from D.S. D.S. secured funding and supervised the project.

## Acknowledgements

Codex (OpenAI) was used to assist with codebase production, organization, and refactoring. ChatGPT was used for copyediting the manuscript, including improvements to grammar, consistency, and readability. All scientific content and final revisions were produced and approved by the authors.

