## Supplementary Information for "AdaptivePy: a unified Python framework for adaptive sampling in molecular dynamics"

### Supporting Methods

#### Analytic benchmark potentials

The two-dimensional benchmark systems were defined using analytic potential-energy functions $U(x, y)$. The exact functional form and parameter values used for each potential are given below. All coordinates and potential energies are reported in the dimensionless units used for the benchmark simulations.

##### Symmetric four-well potential

The symmetric four-well potential [S1] was defined as a sum of 13 axis-aligned Gaussian terms:

$$U(x,y)=\sum_{g=1}^{13} A_g \exp\left[-\frac{1}{2}\left(\frac{(x-\mu_{x,g})^2}{\sigma_{x,g}^2}+\frac{(y-\mu_{y,g})^2}{\sigma_{y,g}^2}\right)\right]. \tag{S1}$$

The amplitudes were

$$\begin{aligned}(A_1,\ldots,A_{13}) = (&-25,-25,-25,-25,\\ &-50,-50,-50,-50,\\ &-50,-50,-50,-50,60).\end{aligned}$$

The Gaussian centers were

$$\begin{aligned}
(\mu_{x,1},\mu_{y,1}) &= (0.2,1.0), & (\mu_{x,2},\mu_{y,2}) &= (1.0,0.2),\\
(\mu_{x,3},\mu_{y,3}) &= (1.8,1.0), & (\mu_{x,4},\mu_{y,4}) &= (1.0,1.8),\\
(\mu_{x,5},\mu_{y,5}) &= (0.4,1.0), & (\mu_{x,6},\mu_{y,6}) &= (0.8,1.0),\\
(\mu_{x,7},\mu_{y,7}) &= (1.2,1.0), & (\mu_{x,8},\mu_{y,8}) &= (1.6,1.0),\\
(\mu_{x,9},\mu_{y,9}) &= (1.0,0.4), & (\mu_{x,10},\mu_{y,10}) &= (1.0,0.8),\\
(\mu_{x,11},\mu_{y,11}) &= (1.0,1.2), & (\mu_{x,12},\mu_{y,12}) &= (1.0,1.6),\\
(\mu_{x,13},\mu_{y,13}) &= (1.0,1.0).
\end{aligned}$$

The coordinate-wise variances were

$$\begin{aligned}
(\sigma_{x,g}^2,\sigma_{y,g}^2) &= (0.008,0.008), & g &= 1,\ldots,4,\\
(\sigma_{x,g}^2,\sigma_{y,g}^2) &= (0.04,0.02), & g &= 5,\ldots,8,\\
(\sigma_{x,g}^2,\sigma_{y,g}^2) &= (0.02,0.04), & g &= 9,\ldots,12,\\
(\sigma_{x,13}^2,\sigma_{y,13}^2) &= (0.02,0.02).
\end{aligned}$$

**Müller–Brown potential**

The Müller–Brown potential [S2] was defined as

$$U(x,y)=\sum_{g=1}^{4} A_g \exp\left[a_g(x-x_g)^2+b_g(x-x_g)(y-y_g)+c_g(y-y_g)^2\right]. \tag{S2}$$

The parameters were

$$(A_1, A_2, A_3, A_4) = (-200, -100, -170, 15),$$
$$(a_1, a_2, a_3, a_4) = (-1.0, -1.0, -6.5, 0.7),$$
$$(b_1, b_2, b_3, b_4) = (0.0, 0.0, 11.0, 0.6),$$
$$(c_1, c_2, c_3, c_4) = (-10.0, -10.0, -6.5, 0.7),$$
$$(x_1, x_2, x_3, x_4) = (1.0, 0.0, -0.5, -1.0),$$
$$(y_1, y_2, y_3, y_4) = (0.0, 0.5, 1.5, 1.0).$$

**Wave-split potential**

The wave-split potential [S3] was defined as

$$U(x, y) = A \exp\left\{ -\frac{1}{2}\left[ B\left(1 - \tanh\left(C - |s_x x|\right)\right) + \frac{1}{2}\left( s_y y - \frac{\sin\left(2\pi s_x x / P\right)}{q} \right)^2 \right] \right\}. \tag{S3}$$

The parameter values were

$$A = -80, \quad B = 20, \quad C = 8,$$
$$s_x = 3, \quad s_y = 2, \quad P = 4, \quad q = 0.8.$$

Substitution of these values gives the explicit expression

$$U(x, y) = -80 \exp\left\{ -\frac{1}{2}\left[ 20\left(1 - \tanh\left(8 - |3x|\right)\right) + \frac{1}{2}\left( 2y - \frac{\sin\left(2\pi(3x)/4\right)}{0.8} \right)^2 \right] \right\}. \tag{S4}$$

**Smooth-minima potential**

The smooth-minima potential [S3] was defined as

$$U(x, y) = \sum_{g=1}^{5} A_g \exp\left[ -w_{x,g}(x - x_g)^2 - w_{y,g}(y - y_g)^2 \right]. \tag{S5}$$

The amplitudes were

$$(A_1, A_2, A_3, A_4, A_5) = (-60, -60, -60, -60, -60).$$

The centers were

$$
\begin{aligned}
(x_1, y_1) &= (-1.1, 0.0), \quad & (x_2, y_2) &= (0.8, -0.3), \\
(x_3, y_3) &= (-1.1, -1.2), \quad & (x_4, y_4) &= (1.55, 0.4), \\
(x_5, y_5) &= (0.0, 0.5).
\end{aligned}
$$

The coordinate-specific width parameters were

$$
\begin{aligned}
(w_{x,1}, w_{y,1}) &= (1.0, 1.5), \quad & (w_{x,2}, w_{y,2}) &= (2.0, 2.0), \\
(w_{x,3}, w_{y,3}) &= (6.0, 3.5), \quad & (w_{x,4}, w_{y,4}) &= (5.0, 5.0), \\
(w_{x,5}, w_{y,5}) &= (5.0, 5.0).
\end{aligned}
$$

**Triple-well potential**

The triple-well potential [S4] was defined as

$$
\begin{aligned}
U(x, y) = 3 \exp &\left[ -x^2 - \left( y - \frac{1}{3} \right)^2 \right] \\
&- 3 \exp \left[ -x^2 - \left( y - \frac{5}{3} \right)^2 \right] \\
&- 5 \exp \left[ -(x-1)^2 - y^2 \right] \\
&- 5 \exp \left[ -(x+1)^2 - y^2 \right] \\
&+ 0.2 x^4 + 0.2 \left( y - \frac{1}{3} \right)^4 .
\end{aligned}
\tag{S6}
$$
